# Seasonal dynamics of the wild rodent faecal virome

**DOI:** 10.1101/2022.02.09.479684

**Authors:** Jayna Raghwani, Christina L. Faust, Sarah François, Dung Nguyen, Kirsty Marsh, Aura Raulo, Sarah C. Hill, Kris V. Parag, Peter Simmonds, Sarah C. L. Knowles, Oliver G. Pybus

## Abstract

Viral discovery studies in wild animals often rely on cross-sectional surveys at a single time point. As a result, our understanding of the temporal stability of wild animal viromes remains poorly resolved. While studies of single host-virus systems indicate that host and environmental factors influence seasonal virus transmission dynamics, comparable insights for whole viral communities in multiple hosts are lacking. Leveraging non-invasive faecal samples from a long-term wild rodent study, we characterised viral communities of three common European rodent species (*Apodemus sylvaticus, A. flavicollis*, and *M. glareolus*) living in temperate woodland over a single year. Our findings indicate that a substantial fraction of the rodent virome is seasonally transient and associated with vertebrate or bacteria hosts. Further analyses of one of the most abundant virus families, Picornaviruses, show pronounced temporal changes in viral richness and diversity, which were associated with concurrent and up to ∼3-month lags in host density, ambient temperature, rainfall and humidity, suggesting complex feedbacks from the host and environmental factors on virus transmission and shedding in seasonal habitats. Overall, this study emphasizes the importance of understanding the seasonal dynamics of wild animal viromes in order to better predict and mitigate zoonotic risks.

## INTRODUCTION

Our knowledge of the global virosphere has rapidly expanded (C. X. Li et al., 2015; Roux et al., 2016; Shi et al., 2018; Zhang et al., 2018), mainly due to decreasing costs and increasing efficiency of high-throughput sequencing. However, while it is now relatively straightforward to genetically characterise host viromes and discover new virus sequences, most studies provide only a glimpse of the circulating virus diversity due to infrequent, non-systematic, and spatially limited sampling of target species. As a result, it is unclear why some viruses are found in some species or populations at specific time points but not in others (Harvey and Holmes, 2022).

While viral discovery studies provide valuable data for understanding the evolutionary history and host range of viruses, they offer only limited insights into what factors shape wild animal viromes. In order to understand viral dynamics in wild populations, we need to move from descriptive host-virus associations to a mechanistic understanding of where and when viruses are transmitted and how entire viral communities (viromes) are shaped by the environment and local host communities (Bergner et al., 2019; Fearon and Tibbetts, 2021). For example, there is compelling evidence that the number of parasites (i.e., richness) in wild animals is strongly influenced by habitat loss and fragmentation (Mbora and McPeek, 2009; Morand et al., 2019), indicating that anthropogenic-mediated changes in host community structures can directly impact parasite community compositions. These findings further highlight why studying community traits, such as parasite richness, is critical for understanding and forecasting zoonotic risk over time and space.

Current knowledge about what factors shape viral communities in wildlife comes from a small but growing number of studies. A comparison of viromes from three parasitic wasp species reared in laboratory conditions suggests that host phylogeny influences viral community structure (Leigh et al., 2018). However, it is unclear whether viromes in wild animals are also commonly predicted by host evolutionary history. Indeed, a study of multiple wild waterbird species sharing habitats found discordance between the host phylogeny and virome composition (Wille et al., 2019). This finding suggests that interspecific interactions and cross-species transmission might break down host phylogenetic structuring of viral communities in wild settings. However, as investigations into community dynamics of viruses in multi-host systems are limited, it remains uncertain how patterns of phylogenetic structuring vary across host and viral taxa, or ecological contexts. Virome composition can also vary among individuals within a species due to demographic and environmental characteristics. For instance, a survey of 24 vampire bat colonies found that virus richness was positively associated with younger age structure, lower elevation, and increasing anthropogenic influence (Bergner et al., 2019). A study on shorebirds also similarly found higher viral richness in younger individuals (Wille et al., 2021), suggesting age structure is a critical determinant of virus diversity in wild animals.

Although evidence suggests that environmental and host factors influence viral communities in wild animals, these surveys have predominantly been cross-sectional, with any given population sampled at a single time point. As a result, we have a sparse understanding of temporal dynamics in wild animal viromes. In particular, fundamental questions such as how viral diversity varies over time and what proportion of viruses are only detected intermittently or at certain times of year in seasonal environments remain unaddressed. Seasonally varying environments profoundly affect wild animals, particularly by regulating critical events in their life cycle (e.g., birth and death rates) and behaviour, including the level and types of social interaction taking place. As a result, many factors affecting viral transmission vary seasonally (Altizer et al., 2006), such as recruitment of susceptible individuals, population density, and contact rates. Furthermore, virus survival in the environment can fluctuate throughout the year and influence onward spread. For instance, the environmental persistence of avian influenza viruses is higher at colder temperatures (Brown et al., 2007) Consequently, it is not surprising that zoonotic virus surveillance studies in reservoir populations have consistently observed seasonal variation in virus prevalence in several host species, such as rodents, bats, birds, and racoons (Amman et al., 2012; Fichet-Calvet et al., 2007; George et al., 2011; Hirsch et al., 2016; Páez et al., 2017). However, except for a handful of studies focusing on specific virus families (e.g., paramyxoviruses in bats and influenza viruses in mallard ducks (Latorre-Margalef et al., 2014; Wille et al., 2018)), investigations into temporal variation in virus diversity in wild animals are rare, leaving a significant gap in our knowledge about viral community dynamics in changeable environments. For example, we may expect to see increases in viral richness during an animal’s breeding season, driven by higher (primarily intra-specific) contact rates. Alternatively, viral community richness or composition may respond to seasonal changes in climate, for instance, if ambient conditions affect viral persistence in the external environment (Brown et al., 2007; Sobsey et al., 1988).

Rodents are a significant zoonotic reservoir globally, and Europe has been identified as a hotspot for zoonotic rodent host diversity (Han et al., 2015). Furthermore, viral metagenomic surveys confirm that wild rodents carry a high and diverse viral burden, which includes several viruses closely related to human pathogens (Drexler et al., 2013, 2012; Firth et al., 2014; Kapoor et al., 2013; Phan et al., 2011; Williams et al., 2018; Wu et al., 2018), including coronaviruses(Wang et al., 2020). Therefore, understanding the composition and dynamics of rodent viromes is an important goal that can help shed light on when and where these host communities may pose the greatest risk of zoonotic spillover to humans. However, our understanding of virus diversity in rodents and what shapes variation in rodent viromes within and among sympatric species remains limited.

To address these questions, we leveraged a long-term capture-mark-recapture study of several sympatric rodent species in Wytham woods, Oxfordshire. Specifically, we characterised the viromes of three common resident species, *Apodemus sylvaticus* (wood mouse), *Apodemus flavicollis* (yellow-necked mouse), and *Myodes glareolus* (bank vole). These three species are ubiquitous across Europe, particularly in woodland habitats. They have fast-paced life histories, with females capable of producing multiple litters in her lifespan, which is typically less than one year. To characterise seasonal variation in viral communities, we generated metaviromic data from pooled faecal samples collected longitudinally from each species over a single year. By combining local microclimate and demographic data from the same period, we explored key factors that predict seasonal variation in the wild rodent virome.

## RESULTS

### Virome dynamics in wild rodents

Over a one-year period (January 2017 to January 2018), we characterised viruses in faeces from a total of 133 individual rodents (57 *A. sylvaticus*, 25 *A. flavicollis*, 51 *M. glareolus*). For each of five 2-3 month sampling intervals, we randomly selected up to 13 samples per species to create species and sampling interval-specific pools for metagenomic sequencing (see Methods for further details). This approach resulted in five pools for both *A. sylvaticus* (wood mouse) and *M. glareolus* (bank vole) and three pools for *A. flavicollis* (yellow-necked mouse) which are less abundant at the sampling site.

Of the total quality-filtered and trimmed reads, 6.77% (∼22.7M/335.9M) were taxonomically assigned to known viruses (see Methods). Figure 1 provides an overview of the viruses detected across all rodent species (hereafter ‘Wytham rodents’). Raw virus abundance ranged from 1.06M to 2.88M reads per pooled sample, with median abundances of 2.27M, 1.40M, and 1. 22M, for wood mice, yellow-necked mice and bank voles, respectively, with read abundance varying somewhat among species and throughout the year (Figure 1A). Although the number of individuals per pooled sample varied between 2-13, this was not significantly correlated with the number of virus genera (i.e., viral richness) in each pooled sample (Pearson correlation = 0.1796; p = 0.56). Rarefaction curves further suggest that viral richness is approaching saturation in the Wytham rodents (Figure S1), indicating additional sampling is unlikely to reveal many more viral genera.

**Figure 1:**
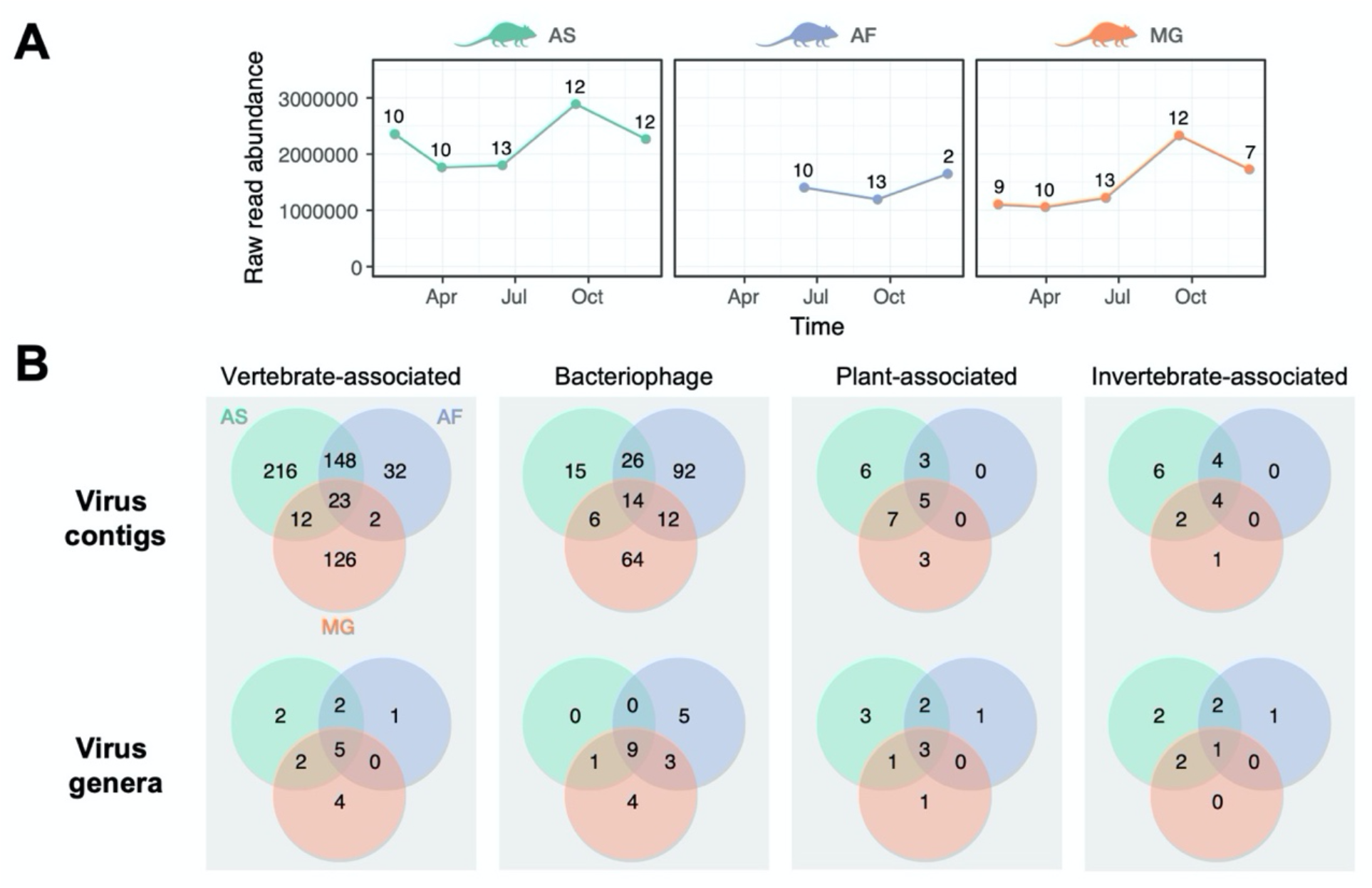
Summary of viral reads detected in the wild rodent faecal virome in Wytham woods over a single year. A) Raw viral read abundance through time for the three host species (AS = wood mouse, AF = yellow-neck mouse, and MG = bank vole). Numbers above each point indicate the number of individuals included in each pooled sample. Timepoints correspond to the midpoint of the sampling interval (see main text). B) Taxonomic assignment of viral reads by contigs and genera across four main groups (based on minimum contig size of 200 nucleotides and applying a minimum threshold abundance of 1 read per million in each pooled sample and restricting to contigs with at least 20 reads). Green, blue, and orange corresponds to AS, AF, and MG, respectively.

The majority of virus contigs are associated with virus families that infect vertebrates or bacteria (Figure 1B). This pattern is somewhat unexpected as the viral enrichment protocol used in this study was optimised for characterising viral RNA in encapsulated viruses (see Methods), regardless of their host association. Specifically, the frequency of bacteriophage contigs is notable since most bacteriophages have double- or single-stranded DNA genomes, although our protocols should also detect DNA viruses undergoing active replication or transcription. Alternatively, bacteriophages could be preferentially enriched in shotgun metagenomic datasets due to their large genome sizes (>100kb). While this might be a contributing factor, the most common bacteriophages in the Wytham rodent virome belonged to *Leviviridae* (+ssRNA) and *Microviridae* (ssDNA), which have genome sizes ranging from 4 to 6.5kb.

While a substantial proportion of contigs were host species-specific (27.5% in wood mice, 14.4 % in yellow-neck mice, and 25.2% in bank voles), most vertebrate-associated viral contigs were detected in at least two host species (Figure 1B). Furthermore, a larger proportion of viral contigs (∼27%) were shared between the two closely related mouse species (wood mouse and yellow-necked mouse) than between mice and voles (13%). Altogether, these observations suggest that both host phylogenetic relatedness and sympatry influence virus distribution.

To understand how virus detection varies across the year, we quantified the proportion of viral genera by the number of times it was detected (Figure 2A) and in each sampling interval (Figure 2B). Overall, 62.4% (116/186) of viruses were only observed in one or two intervals. A similar trend was noted for both vertebrate-associated viruses (19/38 = 50%) and bacteriophage (41/65 = 63.1%), indicating that most viruses in wild rodents are observed intermittently. The number of detected viruses varied seasonally, with the highest percentage observed in the third sampling period (Figure 2B), which corresponds to spring/summer months when population density is low. Furthermore, most viruses (74%) were detected between the third and last sampling periods, i.e., the spring/summer and autumn/winter months (Figure 2B).

**Figure 2:**
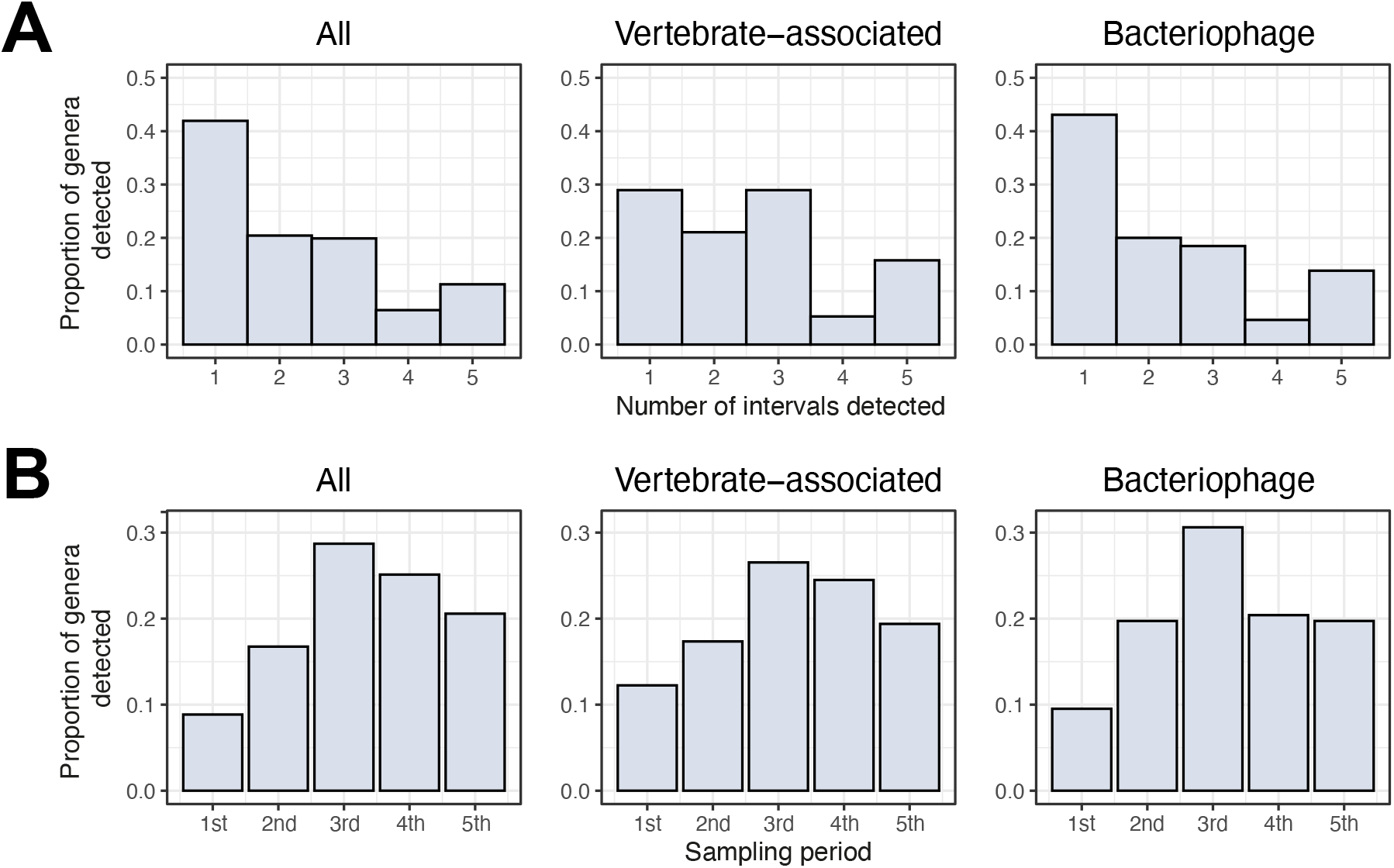
Variation in virus detection across the year. A) Histogram showing the proportion of viral genera by the number of times it was detected across the five sampling intervals for all viruses, vertebrate-associated viruses, and bacteriophages. B) Histogram summarising the proportion of viral genera detected in each sampling period (1 = Jan-Feb 2017, 2 = Mar-Apr 2017, 3 = May-Jul 2017, 4) Aug-Oct 2017, 5) Nov-Jan 2017/18) for all viruses, vertebrate-associated viruses, and bacteriophage.

As sample pooling is likely to conceal viruses at low prevalence or viraemia, variation in virus detection (Figure 2) will reflect changes in both virus abundance and occurrence. We used additive diversity partitioning (Crist et al., 2003) to evaluate how much variation in viral diversity was observed at different levels – within a (pooled) sample, between samples and between sampling periods (Table 1). When considering viromes of all three host species together, around a third of viral diversity (32%) was observed within pooled samples, 21% was observed between pooled samples within a given sampling period (i.e. among species, Table 1), while nearly half of all viral diversity (47%) arose between sampling periods. In both wood mice and bank voles, approximately equal proportions of viral richness occurred within samples (∼48%) and between sampling periods (51-52%). However, in yellow-necked mice, the proportion of virus richness arising across sampling periods was lower (37%) than the other host species (Table 1). This difference is likely to reflect sampling bias, particularly as faecal samples from yellow-necked mice were only available for three out of the five sampling periods. Nevertheless, these findings suggest a significant turnover in viral diversity through time in Wytham rodents and that the structure of viral communities in wild rodents is highly transient.

**Table 1.** Hierarchical partitioning of total virus richness.

| Host group | Mean virus richness | Level | % |
| --- | --- | --- | --- |
| All | 31.92 | Within sample | 32.25 |
|  | 20.88 | Between samples | 21.1 |
|  | 46.20 | Between sampling periods | 46.67 |
|  | 99 | Total | 100 |
| *A. sylvaticus* | 25.4 | Within sample | 47.92 |
|  | 27.6 | Between sampling periods | 52.1 |
|  | 53 | Total | 100 |
| *A. flavicollis* | 38.33 | Within sample | 62.84 |
|  | 22.67 | Between sampling periods | 37.16 |
|  | 61 | Total | 100 |
| *M. glareolus* | 34.6 | Within sample | 48.73 |
|  | 36.4 | Between sampling periods | 51.27 |
|  | 71 | Total | 100 |

### Extensive circulating virus diversity

Focusing on the vertebrate-associated and bacteriophage virus groups, closer examination of the temporal patterns confirmed that considerable virus diversity was detected in the Wytham rodents, corresponding to different virus families, genera, and genome architectures displaying highly variable patterns of seasonal detection (Figure 3). The most prevalent viruses belong to *Picobirnaviridae* (vertebrate-associated) and *Leviviridae* (bacteriophage), which were detected throughout the year at high read abundance (ranging from 0.66M-2.32M for *Picobirnaviridae* and 0.29M-1.35M for *Leviviridae* in pooled samples). Other common vertebrate-associated viruses were members of several non-enveloped ssRNA virus families, such as *Picornaviridae, Astroviridae, Hepeviridae*, and dsRNA virus family, *Reoviridae*. We also detected multiple enveloped RNA and reverse-transcribing viruses (*Betaretrovirus, Betacoronavirus*, an unclassified *Paramyxovirus*, and a *Torovirus*) in yellow-necked mice and bank voles.

**Figure 3:**
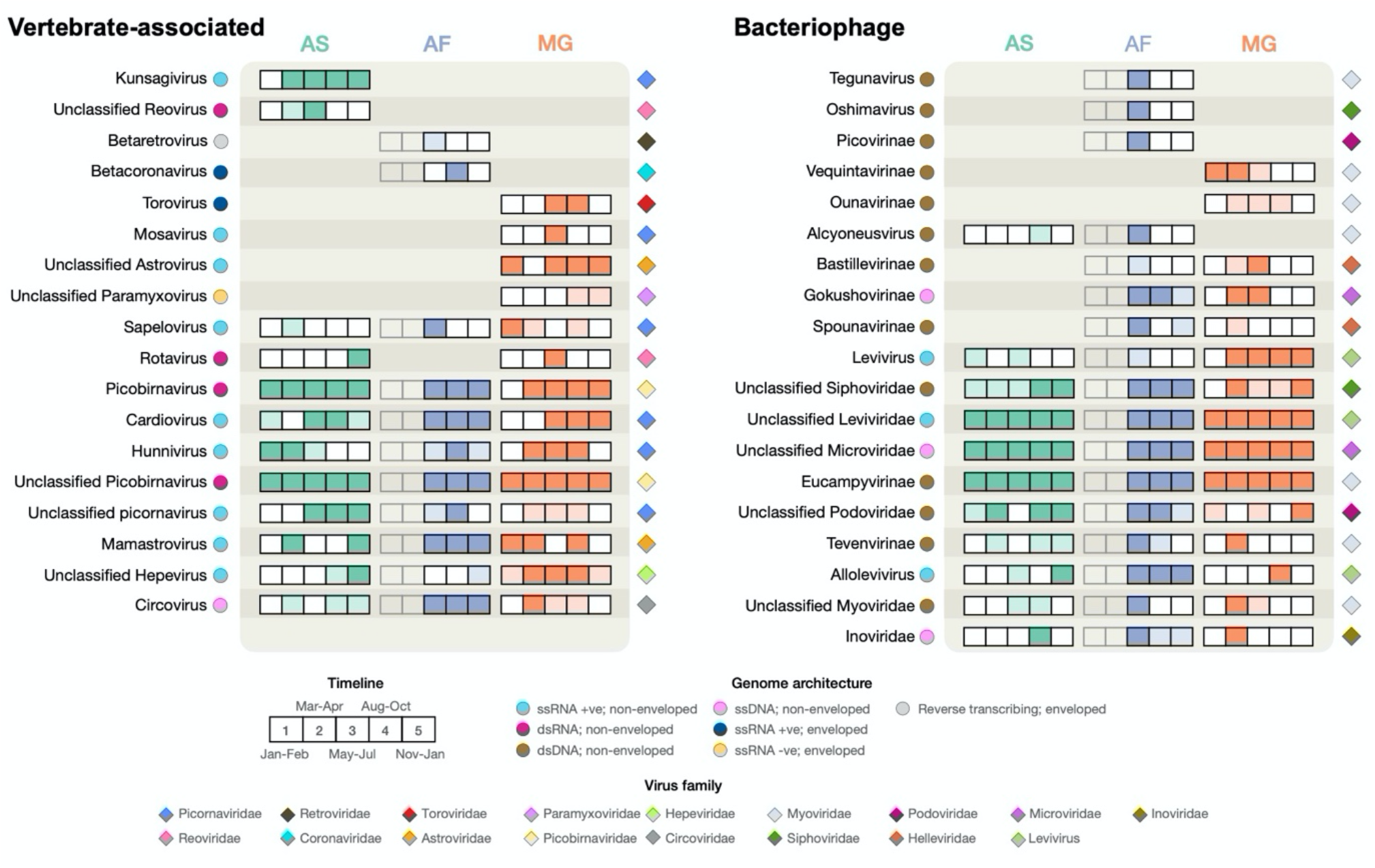
Temporal patterns of vertebrate-associated and bacteriophage virus genera in Wytham rodents. Each box within a set of five corresponds to a distinct sampling interval (five intervals for AS and MG and three for AF). Solid filled boxes indicate that at least 20 reads were detected for a virus genus in a particular host, while light shaded boxes indicate less than 20 reads, and white boxes indicate no reads were detected. Coloured circles indicate the virus (enveloped or non-enveloped) and genome architecture, and coloured diamonds indicate the virus family.

Apart from picornaviruses, more resolved taxonomic classification of the most common vertebrate-associated and bacteriophage virus families, e.g., *Picobirnaviridae, Leviviridae*, and *Microviridae*, was not possible due to poor representation of these taxa in reference databases. As a result, it is challenging to ascertain more detailed information about these viruses. For example, which hosts do the bacteriophage infect and how many distinct virus species (i.e., virus genomes) are there? We aimed to partly address the latter by considering virus contigs that are similar in length to complete genomes (see Table S1), many of which are likely to represent new viruses. Based on this simple approach, there appear to be potentially 114 putative *Picobirnavirus* genomes (which are bisegmented), 21 putative *Levivirus* genomes, and nine putative *Microvirus* genomes (Table S1).

### Seasonal co-circulation of picornaviruses

In Wytham rodents, picornaviruses were the most common and taxonomically well-characterised viruses. Furthermore, as they contain several important pathogens that affect human and animal health (e.g., *Enterovirus* and *Apthovirus*), we undertook a more detailed analysis to understand seasonal variation in picornavirus abundance and diversity. We assembled eight virus contigs for the most prevalent picornaviruses (see Table S2 for further details), representing partial and near-complete genomes. The eight genome sequences correspond to six distinct genera (Figure 4A) and share between 48 and 95% amino acid sequence identity with their closest BLAST hit, which were primarily associated with mammalian hosts, such as bats and other rodent species. The normalised read abundance (the number of reads divided by the number of individuals included in the pooled sample) showed strong seasonal variation for all six viruses (Figure 4B). Furthermore, the seasonal patterns of detectability and peak abundance varied strikingly across these picornaviruses (Figure 4B). For example, *Mosavirus* and *Sapelovirus* were only observed at a single time point and most abundant in early summer (between May and July), while others (e.g., *Hunnivirus* and *Kunsagivirus*) were detected in multiple consecutive periods and reached peak abundance in late summer (between August and October). Importantly, this suggests that even for related viruses, there may be marked variation in the underlying drivers of transmission. Our data also demonstrated that while multiple distinct picornaviruses co-circulate in all three rodent species, for the most part, these viruses are disproportionately associated with a single host species (Figure 4B). In particular, the two distinct genome sequences of the “unclassified picornavirus” genus (Figure 4A) were exclusively found in wood mice or yellow-necked mice, but not both, even when these genomes were detected at the same sampling interval (Figure 4B). However, as the yellow-necked mice are less abundant than the other two species, sampling bias and the pooling of samples are likely to affect the observed virus sharing among host species.

**Figure 4:**
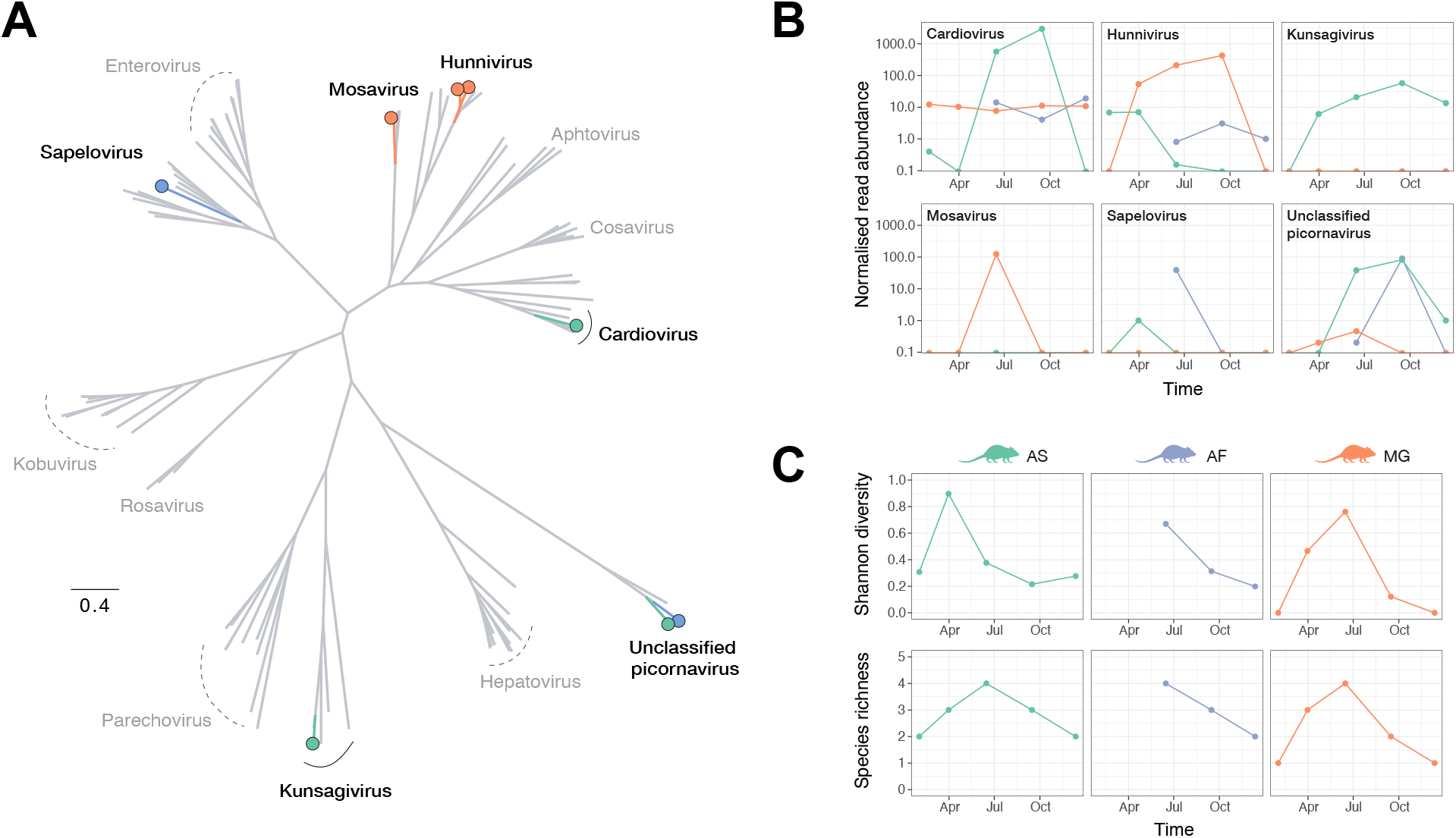
Picornavirus diversity and abundance. A) Evolutionary relationship of picornavirus assembled genomes identified in Wytham rodent faecal virome (coloured by their predominant host association) and a subset of known mammalian picornaviruses (in grey). B) Normalised read abundance of six picornaviruses over time. Colours indicate association with host species (green, orange, and blue correspond to AS, MG, and AF). C) Diversity of picornaviruses, measured as Shannon diversity and richness, over time.

Next, we investigated temporal patterns of picornavirus diversity (Figure 4C). Similar to virus abundance, Shannon diversity and species richness exhibited seasonal variation. The trends are broadly consistent among the three rodent species, with the highest picornavirus diversity occurring between May and July in early summer. However, Shannon diversity notably peaked earlier in wood mice between March and April during the spring months. The pattern is likely driven by the presence of multiple picornaviruses in the wood mice that are at similar low abundance, resulting in high Shannon diversity. As the *Cardiovirus* becomes predominant in the population at later time points, this leads to a concurrent decrease in Shannon diversity as the relative frequencies among co-circulating picornaviruses become unequal. A similar observation is observed in bank voles, where a notable reduction in Shannon diversity in late summer (August to October) coincides with a peak in *Hunnivirus* abundance.

Although we could not examine other common virus families (e.g., *Picobirnaviridae, Leviviridae* and *Microviridae*) detected in the Wytham rodents at the same level of detail due to limited characterisation of virus diversity below the family level, read abundance patterns in wood mice and bank vole provides some insights into their epidemiology (Figure S2). *Picobirnaviridae* and *Leviviridae* showed similar dynamics in both host species and were present at high abundance throughout the year, possibly indicative of a shared transmission mechanism. Conversely, although *Microviridae* abundance exhibited similar dynamics to Picobirnaviridae and Leviviridae in the wood mice, this was not the case in the bank voles (Figure S2). In contrast, changes in *Picornaviridae* abundance in wood mice and bank voles corresponded closely (Figure S2). Overall, these patterns indicate considerable variation in rodent virus transmission with co-circulating viruses characterised by divergent epidemiological dynamics both within and between species.

### Drivers of picornavirus diversity

To explore the predictors of picornavirus diversity in Wytham woods, we focused on wood mice and bank voles, which were sampled in each of the five intervals. We evaluated three environmental variables (temperature, humidity, and rain), using data collected from June 2016 to January 2017 from two weather stations located within the woodlands, together with approximately fortnightly estimates of host population density, calculated as the minimum number known alive (MNKA) per hectare from trapping data. Time series data on picornavirus diversity and the four variables was reconciled using interpolation techniques (see Methods). Specifically, we used a fortnightly interval to derive estimates of all variables at the resolution available for the host density data (Figure S3). We undertook a cross-correlation analysis to select the single most informative time-lag for each of the four variables (temperature, humidity, rain, and host population density), as identified by the highest correlation coefficient (Table S3). The maximum time lag was set as 14 weeks to reflect the expected average lifespan of wood mice and bank voles (∼3 months). This analysis found a notable correlation between picornavirus diversity and the ecological conditions experienced by host species in the preceding weeks and months (Table S3). For Shannon diversity, time lags in the four variables ranged from 10 to 14 weeks in wood mice and 6 to 14 weeks in bank voles, while for species richness, time lags ranged from 2 to 10 weeks for both species (Table S3).

We constructed generalised linear models (GLMs) containing each time-lagged variable as predictors for each host species and diversity metric. Sets of GLMs were reduced accordingly to exclude highly correlated variables (i.e., >0.7; see Table S4). The results indicated that drivers of picornavirus diversity varied by species and diversity metric (Figure 5). For both species, host density in the preceding 2-3 months was negatively correlated with Shannon diversity (Figure 5, Table S5). This suggests that peaks in Shannon diversity followed periods of low population density (in late winter), when populations comprise mostly overwintering individuals, and when home ranges are largest and most overlapping. Shannon diversity was associated with lower temperatures three months previously in wood mice, while Shannon diversity was negatively associated with concomitant humidity in bank voles (Figure 5, Table S5). Despite observing similar trends in viral richness, we found different sets of predictors associated with this metric in the two host species. Rainfall at 10-week and 12-week lags were negatively associated with viral richness for wood mice and bank voles, respectively (Figure 5, Table S5), indicating periods with higher rainfall are generally not conducive for virus survival and/or transmission. We further noted that viral richness was negatively correlated with concomitant humidity in the wood mice. For bank voles, we found that temperature 4-weeks prior and current host density were strongly positively correlated with viral richness (Figures 5; Table S5).

**Figure 5:**
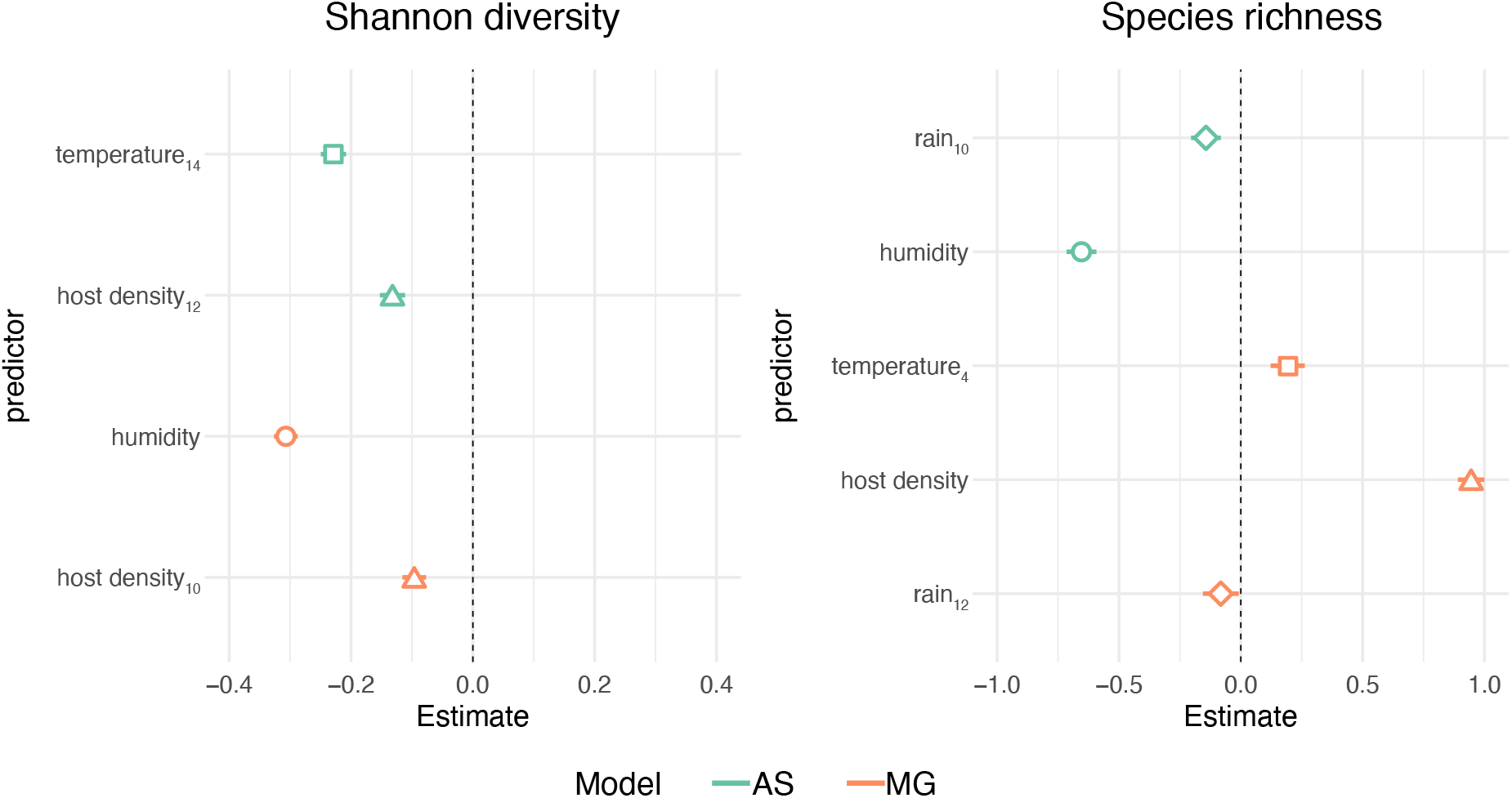
Predictors of picornavirus diversity. Standardised coefficients from the best fit models (mean-centred and scaled by one standard deviation) are illustrated for each diversity metric and species. Subscripts in variable names indicate time lag in weeks. AS = wood mice, MG = bank voles.

Although we found similar predictors associated with Shannon diversity and viral richness in both host species (host density and rainfall at 2-3 months lag, respectively), interspecific differences should be interpreted with caution as the same sets of predictors were not evaluated in each GLM. Consequently, the absence of a predictor in our analyses does not necessarily mean it does not impact viral diversity. To better understand the extent of interspecific variation in shaping virus diversity, additional field data will be required to characterise viral communities on finer temporal scales (e.g., fortnightly or monthly)).

## DISCUSSION

We examined the seasonal dynamics of the faecal virome in three wild rodent species widespread in the UK and Europe. Strikingly, we found extensive virus diversity circulating in these rodents throughout the year. Detected viruses were predominantly associated with vertebrate or bacteria hosts and represented a broad range of virus genomic organisation (RNA and DNA, single- and double-stranded), and virome diversity and community composition varied markedly throughout the year. Although viruses appear to be largely host-specific at the inferred species level, we saw substantial virus sharing among species, particularly among the wood mice and yellow-necked mice, indicating host phylogenetic relatedness is an important determinant of virus ecology. Furthermore, temporal patterns in virus abundance suggest marked variation in the epidemiology of co-circulating viruses, which can differ within and between species. Lastly, seasonal trends in picornavirus diversity suggest that these viral communities are shaped by biological and ecological processes, which likely influence within-host viral dynamics, environmental persistence, and between-host viral transmission.

Our study corroborates previous findings that rodents harbour a substantial and diverse virus burden in the gastrointestinal tract (Firth et al., 2014; Phan et al., 2011; Williams et al., 2018; Wu et al., 2018) with individuals likely encountering a shifting array of seasonally abundant viruses over their lifetimes (Abolins et al., 2018; Firth et al., 2014), contributing to their highly activated immune state (Abolins et al., 2017). Most vertebrate-associated virus genera identified in the Wytham rodents have been detected previously in wild rodents in the USA and China (Firth et al., 2014; Phan et al., 2011; Williams et al., 2018; Wu et al., 2018). *Cardiovirus* and *Picobirnavirus* have been reported in four major untargeted viral metagenomic surveys of wild rodents previously (Firth et al., 2014; Phan et al., 2011; Williams et al., 2018; Wu et al., 2018), suggesting these viruses are widespread and endemic in rodents. Two virus genera detected here have not been reported from wild rodents previously - *Kunsagivirus* (family *Picornaviridae*) and *Torovirus* (family *Tobaniviridae*). Although information about Kunsagivirus is limited (currently only six sequences available in Genbank), *Torovirus* is an enveloped virus commonly found in mammals, including humans, with gastroenteritis (Horzinek et al., 1987; Jamieson et al., 1998). Furthermore, the lower abundance of enveloped viruses than their non-enveloped counterparts is not surprising given their increased lability in the gastrointestinal tract. Though detailed characterisation of enveloped viruses in Wytham rodents was limited, contigs of *Paramyxovirus* detected in bank voles in this study closely matched another *Paramyxovirus* (genus *Jeilongvirus*) isolated from bank voles in Slovenia (Vanmechelen et al., 2018).

There was notable variation in observing a specific virus genus in the Wytham rodents across the year. Some viral genera from the most abundant virus families (*Picobirnaviridae* (vertebrate-associated), *Leviviridae* (bacteriophage) and *Microviridae* (bacteriophage)) were observed at all sampling intervals at high levels in all three species, indicating that they (or their bacterial hosts) persist in the population by establishing a chronic infection or environmental persistence which facilitates frequent reinfection. However, a significant fraction of virus diversity (116/186 genera) was only detected in one or two out of five seasonal sampling periods. Hierarchical analysis of virus richness suggests there is substantial turnover in viral diversity in the Wytham rodents, with around half of diversity absent from each sample interval. Although these results suggest that wild rodents may support different virus epidemiological dynamics within a single year, more in-depth investigations will be required to understand the impact of pooling and sampling effort, particularly for viruses with low prevalence or viraemia, which might appear transient despite continuous circulation. Importantly, however, these findings also highlight that cross-sectional surveys will miss a large proportion of circulating virus diversity, even when samples are taken during times of the year when virus diversity is maximal, such as the spring and summer months in this population.

Despite sharing the same seasonal environment, the factors predicting picornavirus diversity differed between wood mice and bank voles. However, these interspecific differences should be interpreted carefully as different combinations of predictors were evaluated for each diversity metric and host species. Rainfall in the previous months predicted lower virus richness in both host species, suggesting wet conditions reduce picornavirus transmission (and/or environmental persistence) and lead to lower viral species circulating in the following months. We also observed two important concurrent predictors - humidity for wood mice and host density for bank voles – suggesting more picornavirus species were shed in conditions with lower humidity (wood mice) or higher population density (bank voles). Host density in the previous 2-3 months was associated with lower Shannon diversity in both species – several mechanisms could explain this pattern. For example, a higher population density could facilitate certain viruses to dominate transmission events through smaller home range sizes and reduced frequency of contacts, or the increase in density could affect competition and alter within-host replication dynamics. Future studies that incorporate more samples collected at higher frequencies could be used to test such hypotheses explicitly.

The widespread distribution of wood mice and bank voles in the UK makes them highly amenable for long-term field studies and have been previously leveraged to understand natural drivers of virus transmission in wildlife populations (Begon et al., 2009; Carslake et al., 2006; Knowles et al., 2012; Telfer et al., 2007, 2002). While these studies have focused on specific DNA viruses that are endemic in these species, they also observed heterogeneity in rodent virus epidemiology, including between years, host species, individuals, and for different viruses (Begon et al., 2009; Carslake et al., 2006; Knowles et al., 2012; Telfer et al., 2007, 2002).

We detected long time-lags (∼3 months) between some environmental variables and picornavirus diversity, particularly for Shannon diversity. This observation could be because pooled samples were from a time window, where individual samples from 2 to 3 months were aggregated into one ‘timepoint’. Although such temporal pooling is not ideal for time series evaluation, it provides a valid first approximation of important seasonal correlates of viromes and an improvement on previous cross-sectional surveys. We expect many of these viruses to be transmitted between conspecifics through close contacts and between species via the environment. However, the ability for viruses to remain transmissible in the environment is highly variable across taxa. For example, hepatitis A virus (genus *Hepatovirus*, family *Picornavidae*) is very stable under a broad range of temperature, humidity, and pH conditions and can survive over three months in the environment (Sobsey et al., 1988). In contrast, other picornaviruses, such as Foot and Mouth disease virus (genus *Aphthovirus)*, appear to be less stable in the environment, with longer survival times observed at higher humidity and moderate temperatures (Abad et al., 1994; Mbithi et al., 1991; Mielke and Garabed, 2020). Although we observed clear seasonality in picornavirus detection and abundance, given the substantial temporal turnover in viral diversity, it is reasonable to assume that other viruses in Wytham rodents also circulated seasonally, especially those detected transiently in the population (e.g., *Coronavirus, Paramyxovirus*). In the future, using field studies with a higher temporal resolution, we plan to develop mechanistic transmission models in these systems. Mechanistic models could be adapted to other rodent systems to forecast peaks and troughs in epizootics and test potential interventions in settings where zoonotic viruses are a risk to human populations.

Understanding viral community dynamics is key to predicting and mitigating human risk from known and unknown rodent zoonoses. Improvements in sequencing technology that enable identification and monitoring of RNA viruses longitudinally in wildlife are crucial to establishing the spatial, temporal and environmental factors that determine zoonotic risk. Previous work has shown that specific rodent-borne zoonotic viruses exhibit strong seasonal dynamics in the reservoir population (Fichet-Calvet et al., 2007; Luis et al., 2015; Tian et al., 2017). Nevertheless, by quantifying the virome, we can identify the co-occurrence of a community of viruses, their transmission across the year, and associations with the environment and host ecology. This step moves forward our current knowledge about the seasonal dynamics of viral communities and contributes to a more comprehensive understanding of virus transmission ecology in wildlife populations.

## METHODS

### Study population

Wild rodents were trapped and sampled over a one-year period (January 2017 to January 2018) in Wytham Woods (51° 46’N,1°20’W), a 385-ha mixed deciduous woodland near Oxford, UK. Three common rodent species are regularly caught at this site: two species of *Apodemus* mice (*Apodemus sylvaticus* and A. *flavicollis*, with *A. sylvaticus* more abundant) and the bank vole (*Myodes glareolus*). These are non-group-living, omnivorous woodland rodents with overlapping home ranges that show seasonal variation in reproduction, mortality, diet (Watts, 1968) and social interactions (Raulo et al., 2021). One night of trapping on a single c. 2.4ha trapping grid was carried out approximately fortnightly year-round. Small Sherman traps (baited with six peanuts, a slice of apple, and sterile cotton wool for bedding material) were set at dusk and collected at dawn the following day. Newly captured individuals were PIT-tagged for unique identification. Faecal samples were collected from the bedding material with sterilized tweezers and frozen at -80°C within 10 hours of trap collection. Traps that showed any sign of animal contact (traps that held captured animals and trigger failures where an animal has interfered with the trap but not been captured) were washed thoroughly with bleach in between trapping sessions to prevent cross-contamination. All live-trapping work was conducted with institutional ethical approval and under Home Office licence PPL-I4C48848E.

### Sample selection and processing

We randomly selected 133 individual faecal samples (57 *A. sylvaticus*, 25 *A. flavicollis*, 51 *M. glareolus*). Five sampling intervals were defined, which took into account the breeding cycle of the three rodent species. These were 1) Jan-Feb 2017, 2) Mar-Apr 2017, 3) May-Jul 2017, 4) Aug-Oct 2017, 5) Nov-Jan 2017/18. Samples were pooled by species and sampling interval, using equal aliquots of 40mg faeces per individual per pool. For the last sampling interval where there were fewer individuals of *A. flavicollis* and *M. glareolus* available (2 and 7 respectively), larger volumes were used for pooling (150mg and 70 mg respectively) to ensure sufficient material for sequencing.

The samples were processed as follows to enrich for RNA within encapsulated viruses: 1) faecal supernatants from pooled samples were filtered through a 0.45nm pore filter; 2) RNase treatment (RNase One) to remove non-encapsulated RNA from sample; 3) RNA extraction using Zymo Quick Viral RNA and RNA Clean and Concentrator 5 kits; 4) DNA digestion following RNA extraction; 5) ribosomal depletion during sequencing library preparation. Sequencing library preparation, which included cDNA synthesis, and sequencing was carried out by the Oxford Genomics Centre on Illumina NovaSeq 6000 platform.

### Viral genomes reconstruction

A total of 352,872,111 pair-end reads of 150 base-pairs (bp) were obtained after sequencing. Illumina adaptors were removed, and reads were filtered for ≥q30 quality and read length >45bp) using cutadapt 1.18 (Martin, 2011). A total of 335,917,017 cleaned reads were *de novo* assembled into contigs by MEGAHIT 1.2.8 with default parameters (D. Li et al., 2015). Taxonomic assignment was achieved on contigs through searches against the NCBI RefSeq viral database using DIAMOND 0.9.22 with an e-value cutoff of <10^−5^ (Buchfink et al., 2014). All contigs that matched virus sequences were selected and used as queries to perform reciprocal searches on NCBI non-redundant protein sequence database with an e-value cutoff of <10^−5^ to eliminate false positives(Altschul et al., 1990). Viral sequences were classified as viral operational taxonomic units (vOTU). vOTU contigs completion and coverage was assessed by iterative mapping using BOWTIE2 2.3.4.3 (Langmead, 2010) and BBMap 35.34 (Bushnell, 2014). Open Reading Frames (ORFs) were searched using ORF finder (parameters: minimum ORF size of 300 bp, standard genetic code, and assuming there are start and stop codons outside sequences) on Geneious prime 2019.1.1 (Kearse et al., 2012).

### Virus abundance and diversity

Analyses of virus abundance and diversity was undertaken in R version 4.1.1 (The R Core Team, 2021), and primarily used the tidyverse package for plotting the data (Wickham et al., 2019). To reduce the impact of contamination in our analyses, we excluded viral contigs with less than one read per million. To examine the distribution of viral contigs in the different host species and virus groups, we used the “ggvenn” library (Yan, 2021). We further restricted the data to viral contigs with at least 20 mapped reads for this analysis. The abundance of common vertebrate-associated and bacteriophage viruses in the three species over time was created with Adobe Illustrator 2021. Rarefaction curves, virus diversity, and additive partitioning diversity were calculated using the “vegan” library (Oksanen et al., 2020).

To reconstruct the picornavirus phylogeny, we assembled a multiple protein sequence alignment of 93 whole picornavirus genome sequences from the NCBI RefSeq viral database and eight picornavirus genome sequences identified in this study. Maximum-likelihood phylogeny was inferred with IQ-TREE v. 2.1.3 (Minh et al., 2020) using the best substitutional model identified by ModelFinder (Kalyaanamoorthy et al., 2017)

### Drivers of picornavirus diversity

We evaluated drivers of picornavirus diversity with Gaussian distributed generalised linear models. We focused on wood mice and bank voles, which were sampled throughout the year. We evaluated three environmental variables (temperature, humidity, and rain), using data collected from June 2016 to January 2017 from two microclimate stations within the woodlands, as well as host population density for the respective species, as measured by the minimum number known alive per hectare over the study period. To match the time series data for the different variables, we interpolated temporal changes over the study period using either locally estimated scatterplot smoothing (LOESS) or generalised additive models (GAMs) in R using the “stats” and “mgcv” libraries. Smoothed curves for Shannon diversity, species richness, and host population density were estimated with LOESS, while the smooth curves for the microclimatic data (temperature, humidity, and rain) were inferred with GAMs.

Time-lagged relationships between picornavirus diversity (Shannon diversity and species richness) with the four variables (temperature, humidity, rain, and host population density) were explored per metric and host species by using the cross-correlation function (ccf) in R. We evaluated lags from 0 to 14 weeks, in two weeks increments, for all four variables. Time lags were identified using significant residual auto-correlation values. If multiple lags were identified per variable, we selected the lag with the highest significant residual auto-correlation value (see Table S3). The maximum lag was set at 14 weeks to reflect the average lifespan of the wild rodents in the study (approximately three months).

We considered four separate GLMs per host species and diversity metric (i.e. AS vs Shannon diversity, MG vs Shannon diversity, AS vs species richness, and MG vs species richness) to evaluate drivers of picornavirus. However, prior to undertaking a GLM analysis, correlations among the four variables (with or without lags as determined by the cross-correlation analysis; Table S3) for each metric and host species were visually assessed in each GLM in R using the library “corrplot” (Wei and Simko, 2021). If the correlation coefficient was 0.7 or greater, we reduced the sets of GLMs considered accordingly (Table S4). We used the library “AICmodvg” (Mazerolle, 2020) for model selection, which considers the Akaike Information Criterion (AIC) and the number of parameters to determine the best fit model. Lastly, the GLM results were plotted using the library “jtools” (Long, 2020).

## ACKNOWLEDGEMENTS

The viral metagenomics study was supported by the British Ecological Society small research grant (award no. SR18/1519) awarded to JR. We would like thank Dr Curt Lamberth, The Wytham Estate, for providing the microclimate data. SF and OGP are supported by BBSRC grant BB/T008806/1.s. CLF was funded by a NERC Fellowship (NE/V014730/1), SCLK by a NERC fellowship (NE.L011867/1) and an ERC Starting Grant (MUSMICRO, Project number 851550), KM by an RVC studentship, and AR by a Clarendon Scholarship.

## AUTHOR CONTRIBUTIONS

Conceptualisation: JR; Funding acquisition: JR; Resources: SCH, PS, and OGP; Investigation: JR, CLF, SF, DN, KVP, KM, AR, and SCLK; Visualisation: JR and CLF; Project administration: JR; Supervision: JR; Writing – original draft: JR; Writing – review & editing: JR, CLF, SF, DN, SCH, KVP, KM, AR, PS, SCLK, and OGP.

## DATA ACCESIBILITY STATEMENT

The raw sequencing data generated in this study have been deposited in the Sequence Read Archive (BioProject ID: PRJNA803204) under the accession numbers: SRX14033113-SRX14033125. Code associated with this research is available at https://github.com/jnarag/Wytham-rodent-virome.

