## Supplementary material for "Seasonal dynamics of the wild rodent faecal virome": Table S1, Table S2, Figure S1, Figure S2, Figure S3, Table S3, Table S4, Table S5

**Supplementary Information**

***Table S1:*** *Number of assembled virus contigs that are similar in length to complete virus genomes in the most abundant virus families in observed Wytham rodent virome.*

| **Group** | **Virus family** | **Number of assembled**  **contigs (length)** |
| --- | --- | --- |
| Vertebrate-associated | *Picobirnaviridae* (dsRNA) | 1) 114 (2100-3000bp^a^)  2) 154 (1200-1900bp^b^) |
| Vertebrate-associated | *Picornaviridae* (ss+RNA) | 8 (>4500bp) |
| Bacteriophage | *Leviviridae* (ss+RNA) | 21 (>3500bp) |
| Bacteriophage | *Microviridae* (ssDNA) | 9 (>4000bp) |

^a^ range based on segment 1

^b^ range based on segment 2

***Table S2:*** *Description of Picornavirus genomes assembled in Wytham rodents*

| **Contig name** | **Genus** | **Genome length** | **Complete**  **(Y/N)** |
| --- | --- | --- | --- |
| Wytham cardiovirus | *Cardiovirus* | 8205 bp | Y |
| Wytham mosavirus | *Mosavirus* | 8114 bp | Y |
| Wytham kunsagivirus | *Kunsagivirus A* | 4851 bp | N |
| Wytham hunnivirus (a) | *Hunnivirus* | 7737 bp | Y |
| Wytham hunnivirus (b) | *Hunnivirus* | 7405 bp | Y |
| Wytham sapelovirus | *Sapelovirus* | 7553 bp | Y |
| Unclassified picornavirus 1 (a) | Unclassified picornavirus 1 | 9085 bp | Y |
| Unclassified picornavirus 1 (b) | Unclassified picornavirus 1 | 9217 bp | Y |

***Figure S1:*** *Rarefaction curves for all viruses, vertebrate-associated viruses, and bacteriophage.*

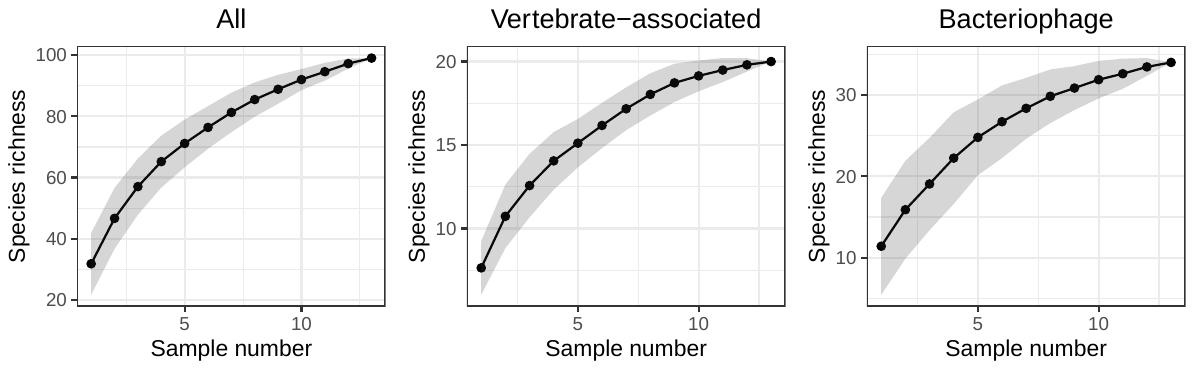

***Figure S2:*** *Normalised read abundance (log-scale) of the most abundant vertebrate-associated and bacteriophage virus families in wood mice and bank voles.*

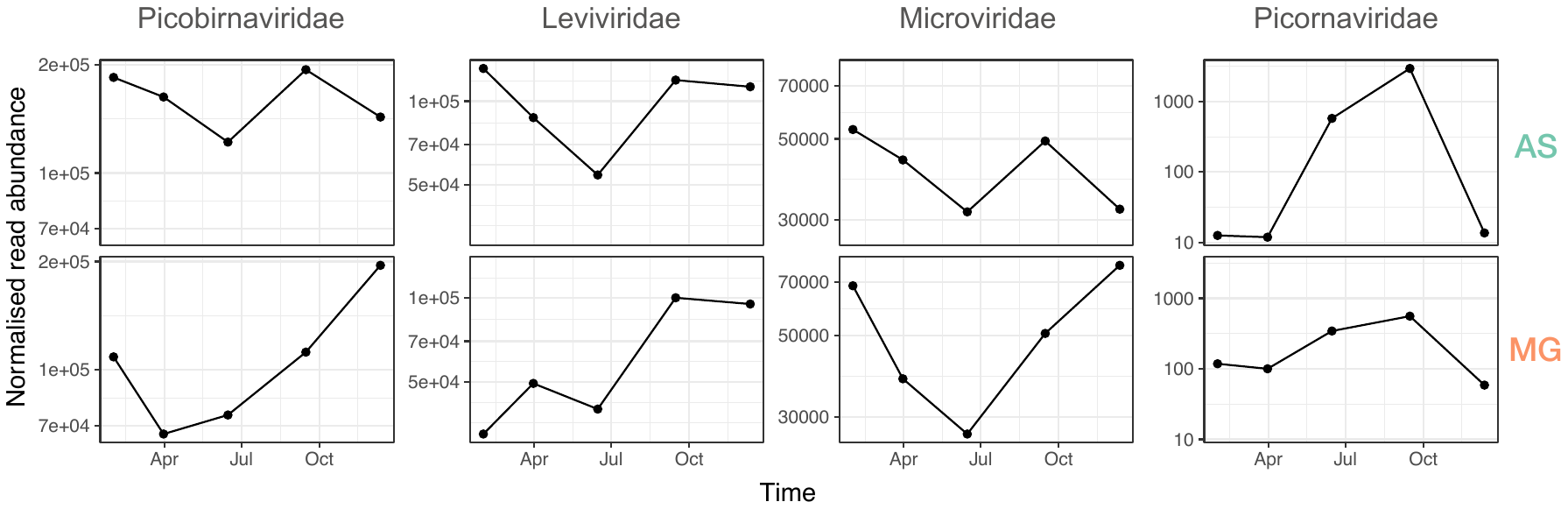

***Figure S3:*** *Summary plot of variables used to explore predictors of picornavirus diversity. A) Shannon diversity and species richness of picornaviruses, together with host population density (measured as MNKA per ha) for wood mice (AS) and bank voles. Open circles indicate imputed values, while filled circles correspond to observed values. B) Daily inferred temperature (mean), humidity (mean), and rainfall (summed per 24 hour period) in Wytham woods using two microclimate stations.*

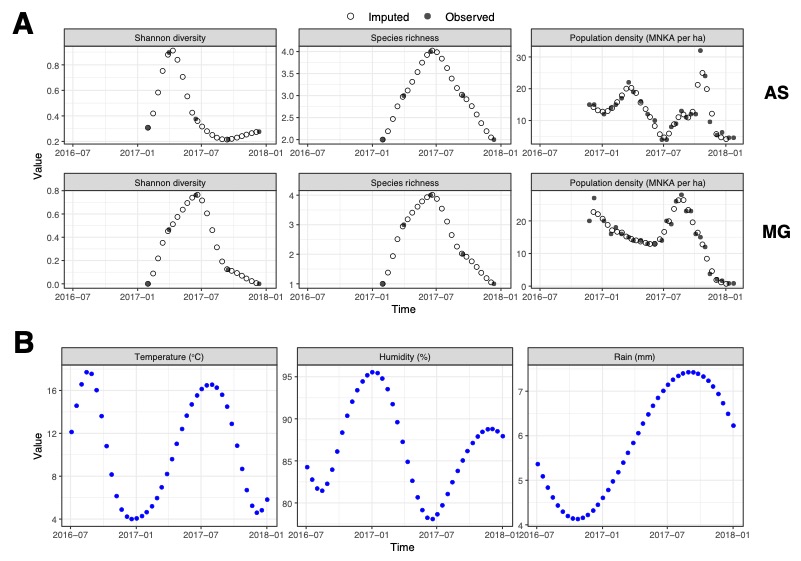

***Table S3:*** *Summary of cross-correlation analysis of four key variables. Lags (in weeks) were identified based on the maximum significant residual auto-correlation values (r_xy_).*

|  | **AS** | | **MG** | |
| --- | --- | --- | --- | --- |
| **Variables** | Shannon diversity  (Lag in weeks; r_xy_) | Species richness  (Lag in weeks; r_xy_) | Shannon diversity  (Lag in weeks; r_xy_) | Species richness  (Lag in weeks; r_xy_) |
| Temperature | 14; 0.64 | 2; 0.83 | 6; 0.74 | 4; 0.77 |
| Humidity | 10; 0.63 | 0; -0.89 | 0; -0.77 | 0; -0.82 |
| Rain | 0; -0.385 | 10; 0.40 | 14; -0.41 | 14; 0.40 |
| Host density | 12; -0.418 | 0; -0.52 | 10; -0.72 | 0; -0.41 |

***Table S4:*** *Summary of GLMs evaluated for each species and diversity metric. Subscripts in the GLMs indicate the time lag of the variable in weeks. AS = wood mice; MG = bank voles.*

| **Host species** | **Virus diversity metric** | **Model** |
| --- | --- | --- |
| AS | Shannon diversity | 1. Temperature + humidity + host density 2. Temperature + rain + host density 3. Temperature + humidity 4. Temperature + host density 5. Temperature + rain 6. Rain + host density 7. Host density + humidity |
| MG | Shannon diversity | 1. Temperature + humidity 2. Humidity + host density 3. Temperature + rain 4. Rain + host density 5. |
| AS | Species richness | 1. Temperature + rain 2. Temperature + host density 3. Humidity + host density 4. Rain + humidity |
| MG | Species richness | 1. Temperature + humidity + rain 2. Temperature + host density + rain 3. Temperature + host density 4. Temperature + rain 5. Temperature + humidity 6. Humidity + rain 7. Rain + host density |

***Table S5:*** *Scaled coefficients of the best fit models.* Subscripts next to predictor variables indicate species and/or time lags (in weeks). *Predictors are mean-centred and scaled by one standard deviation. Significance levels indicated by *** p < 0.001; * p < 0.05*

| **Predictor** | **Shannon diversity (AS)** | **Shannon diversity (MG)** | **Species richness (AS)** | **Species richness (MG)** |
| --- | --- | --- | --- | --- |
| Temperature_14_ | -1.24***  [-1.35, -1.13] |  |  |  |
| Host density_12 (AS)_ | -0.71***  [-0.83, -0.60] |  |  |  |
| Humidity |  | -1.07***  [-1.14, -1.00] | -1.01***  [-1.10, -0.91] |  |
| Host density_10, MG_ |  | -0.34***  [-0.40, -0.27] |  |  |
| Rain_10_ |  |  | -0.22***  [-0.32, -0.13] |  |
| Temperature_4_ |  |  |  | 0.19***  [0.12, 0.26] |
| Host density_0,MG_ |  |  |  | 0.92***  [0.87, 0.98] |
| Rain_12_ |  |  |  | -0.08*  [-0.15, 0.01] |
